# SIMBA: an agentic AI platform for single-molecule multi-dimensional imaging

**DOI:** 10.64898/2026.04.16.719005

**Authors:** Hongjing Mao, Harsh Mauny, Obblivignes KanchanadeviVenkataraman, Caroline Laplante, Dongkuan (DK) Xu, Yang Zhang

## Abstract

Advances in multi-dimensional imaging method and probe developments have brought super-resolution fluorescence microscopy into a functional era. They capture additional single-molecule fluorescence information concurrently with spatial localization, enabling simultaneous identification of molecular species and interrogation of nanoscale environments with rich, high-dimensional imaging information. However, the adoption of multi-dimensional imaging has been hindered by fragmented analysis workflows, complex parameter tuning, and limited integration of advanced computational methods. Here, we introduce an agentic <u>s</u>ingle-molecule <u>m</u>ulti-dimensional <u>b</u>ioimaging <u>A</u>I, referred to as SIMBA, an AI-driven platform that unifies single-molecule localization, spectral processing and deep learning-based denoising within a single agentic and interactive framework. SIMBA incorporates large language model-based agents capable of interpreting user intent, orchestrating analysis pipelines, and dynamically selecting computational tools for automated data processing. We demonstrate that SIMBA enables supports standard single-molecule localization workflow, functional mapping of nanoscale environmental heterogeneity through single-molecule spectral analysis and denoising using developed supervised learning methods. By integrating extensible tool architectures with human language-guided workflows, SIMBA establishes a new paradigm for intelligent microscopy analysis, lowering barriers to multi-dimensional imaging adoption while enabling scalable, reproducible, and adaptive analysis of complex imaging datasets.

## Introduction

Single-molecule localization microscopy (SMLM) has transformed biological imaging by enabling nanometer-scale spatial resolution (<10 nm) far beyond the optical diffraction limit.^1–5^ Traditional SMLM techniques primarily resolve molecular positions, relying on spatial separation or stochastic activation to distinguish emitters. However, many biological questions require simultaneous access to additional molecular dimensions, including identity, chemical state, and local environment.^6–9^ Multi-dimensional SMLM (md-SMLM) has emerged as a class of powerful single-molecule imaging technologies push SMLM beyond mapping spatial patterns.^6,7,10–12^ Rather than focusing solely on spatial localization, md-SMLM expands the measurement space to incorporate additional single-molecule dimensions, including emission spectra^6,8,11,13–17^, fluorescence lifetime^18,19^, polarization^7,20–22^. By encoding these fluorescent properties in new dimensions at the single-molecule level, md-SMLM enables more robust molecular identification, improved multiplexing capacity, and deeper insight into nanoscale interactions among molecules and at distinct local environment. As an versatile example implementation of md-SMLM, spectrally-resolved super-resolution microscopy^11,14–16^ or spectroscopic SMLM^10,13,23–29^ (sSMLM) developed by others and us incorporates a dispersive optical element into the emission path, allowing each localized emitter to be associated with a spectral signature even if the fluorophores have heavy spectral overlap (**Figure 1a**). This capability enables simultaneous multiplexed super-resolution imaging of subcellular organelles^11^ using spectrally overlapping fluorophores (**Figure 1b**) and parallel multiplexed single-molecule tracking^14^. sSMLM also provides a route to functional nanoscale mapping through probing spectral response of environment-sensitive probes at distinct subcellular physiochemical environments such as chemical polarity or hydrophobicity (**Figure 1c**).

**Figure 1.**
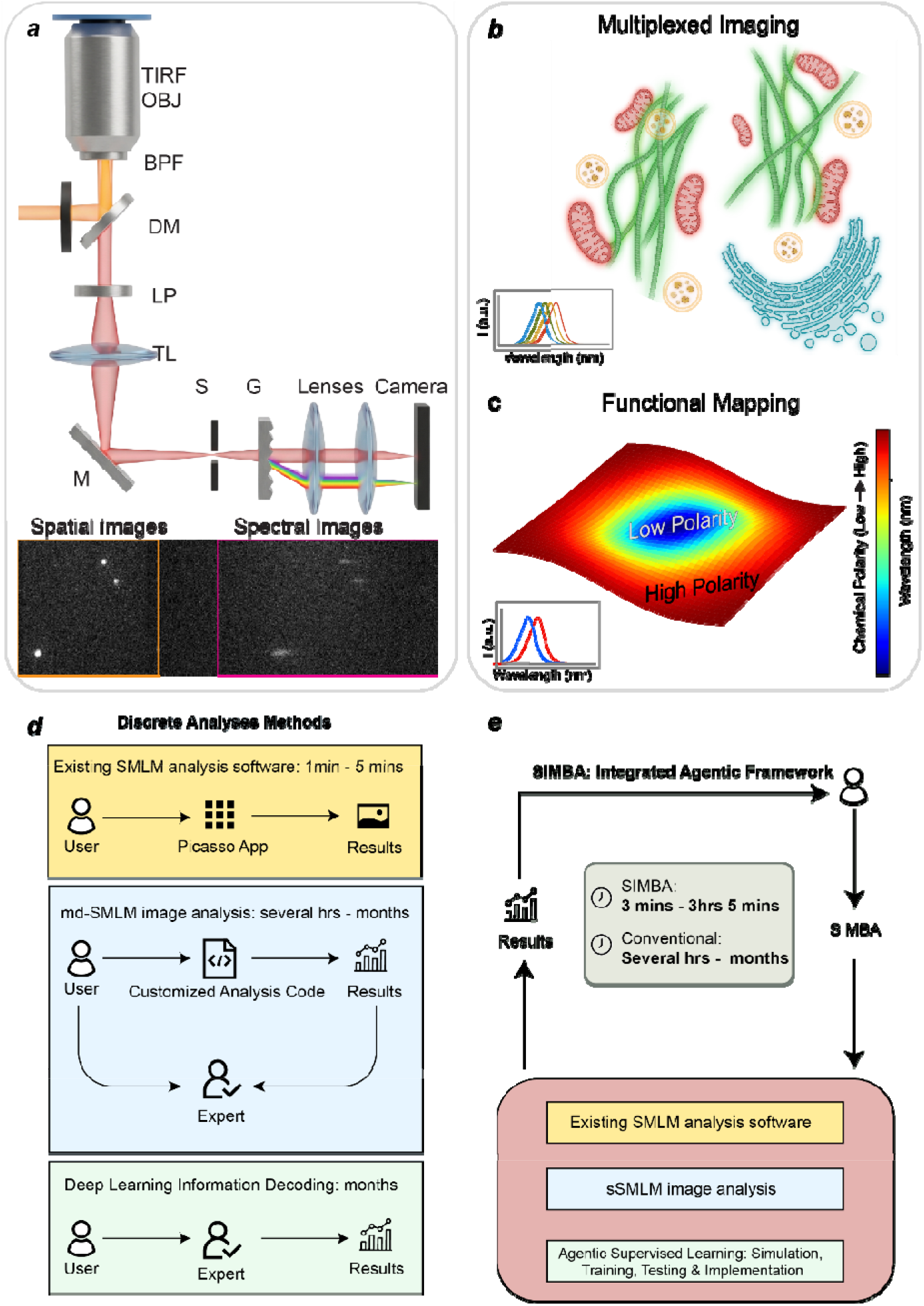
Principles and AI-based agentic analytical framework of md-SMLM. (***a***) Schematic of a spectrally-resolved md-SMLM imaging configuration; (***b***) Demonstration of multiplexed imaging, where distinct spectral signatures enable differentiation of fluorophores labeled with different subcellular organelles (top) with overlappin emission profiles (bottom); (***c***) Illustration of a functional nanoenvironment polarity mapping using polarity-sensitive probes; (***d***) Overview of conventional md-SMLM data analysis workflows, which rely on separate modules; (***e***) Proposed SIMBA framework with a unified automated pipeline, enabling seamless coordination, adaptive workflow orchestration, and natural language guided interactive analysis.

Realizing the full potential of md-SMLM requires tightly integrated workflows that bridge optical system design, fluorophore development, and computational imaging analysis.^10^ While optical implementation and probe engineering are under rapid development, the imaging analysis pipeline is currently underdeveloped. Conventional imaging processing workflows remain fragmented, requiring users to navigate multiple independent tools and programming environments. Typically, the first step in md-SMLM is the standard single-molecule localization process which can be performed using established software such as Python-based Picasso^30^ or Javascript-based ThunderSTORM^31^ as an ImageJ plugin developed for conventional SMLM. Then, it follows by customized information decoding pipelines and, increasingly, separate deep learning models^27,29,29,32–39^ for denoising and signal reconstruction to link the spatial and additional fluorescence patterns. These components are rarely integrated, leading to workflows that depend heavily on manual parameter tuning, iterative troubleshooting, and expert intervention. As a result, analysis is often time intensive, ranging from hours to months, and difficult to reproduce across users and laboratories, as highlighted in **Figure 1d**. Even within a single research group, the development of analysis code is frequently fragmented across projects, contributors, and evolving experimental conditions, resulting in heterogeneous and difficult-to-maintain codebases that further complicate reproducibility and usability. This internal fragmentation can already introduce substantial barriers before considering differences across laboratories. Moreover, deploying such workflows for downstream biomedical researchers, who may not have extensive computational training, remains particularly challenging. The translation from tool development to practical use often requires extensive back-and-forth communication between domain scientists and method developers, significantly slowing the pace of research and limiting broader adoption.

This fragmentation presents a fundamental bottleneck that limits the accessibility and scalability of md-SMLM, particularly for interdisciplinary researchers who may not have specialized computational expertise. We believe that advances in instrument architectures must be co-designed with molecular probes whose photophysical properties are optimized for multi-dimensional encoding, while analysis pipelines must be capable of jointly extracting and interpreting these high-dimensional signals at the single-molecule level. Such end-to-end integration is essential to ensure that gains in one domain, for example spectral resolution or probe sensitivity, are not limited by bottlenecks in another, ultimately enabling robust, scalable, and information-rich md-SMLM measurements.^40^

Recent advances in artificial intelligence (AI), particularly large language models (LLMs)^41–48^, have introduced a new paradigm for interacting with complex computational systems. Unlike conventional automation, which relies on predefined pipelines, agentic AI systems are capable of interpreting user intent, dynamically selecting tools, and orchestrating multi step workflows through iterative reasoning and feedback.^49^ These systems integrate natural language interaction with programmatic execution, enabling users to specify high level goals rather than low-level procedural steps. Emerging evidence across domains suggests that such agent based frameworks can substantially lower barriers to complex data analysis by embedding domain knowledge, adaptive decision making, and interactive refinement within a unified system. Agentic AI systems can interpret natural language instructions, select appropriate computational tools, and iteratively refine analysis strategies. In the context of md-SMLM imaging, agentic AI has the potential to fundamentally reshape how analysis pipelines are designed and used. Rather than treating localization, spectral analysis, and deep learning (DL) as isolated modules (**Figure 1d**), an agentic system can coordinate these components into a cohesive and adaptive workflow. This shift moves the burden of integration and parameter optimization from the user to the system, enabling more efficient exploration of parameter spaces, more consistent analytical outcomes, and more accessible entry points for non experts. Importantly, this paradigm also supports iterative human AI collaboration, where users can guide, inspect, and refine analysis through natural language, while the system maintains computational rigor and reproducibility.

Therefore, we present SIMBA, an agentic AI-driven platform for md-SMLM analysis that unifies single-molecule localization, multi-dimensional fluorescence feature extraction, and advanced DL imaging analytics within a single framework. SIMBA leverages an extensible tool architecture and an LLM-based agent loop to enable human language-guided data processing, automated parameter optimization, and interactive analysis. SIMBA can also be readily extended to excuate localization tasks in conventional SMLM and be transformed to handle other md-SMLM analytics. We demonstrate that SIMBA not only simplifies md-SMLM workflows but also enhances analytical performance and reproducibility, enabling robust multi-target imaging and nanoscale environmental mapping. As shown in **Figgure 1e**, SIMBA incorporates an AI-driven agent loop that interprets user queries, selects appropriate analytical tools, integrates contextual information, and executes workflows end to end. By bridging previously disconnected analysis stages, SIMBA transforms md-SMLM data processing from a fragmented and expert dependent procedure into an automated, interactive, and scalable system. This integration reduces analysis time from hours or months to minutes per dataset, while enabling users to fully leverage the rich information embedded in md-SMLM. This work positions agentic AI as a generalizable framework for next generation scientific infrastructure. By embedding intelligence into the orchestration layer of data analysis, agentic systems can enable new forms of human AI collaboration, accelerate discovery, and democratize access to advanced methodologies across disciplines. SIMBA represents not only a technical advance for md-SMLM, but also a step toward a new paradigm in which complex and interdisciplinary scientific workflows become adaptive, interpretable, and accessible through natural interaction.

## Results

### Computational Implementation of SIMBA

To operationalize this agentic paradigm for md-SMLM, we developed SIMBA as a full stack desktop system that tightly integrates user interaction, agent based orchestration, and domain-specific scientific computing. The overall system design is illustrated in **Figure 2**, which combines a general agent architecture **(Figure 2a**), detailed software implementation (**Figure 2b**), and a domain-specific workflow (**Figure 2c**) tailored for md-SMLM analysis. At the core of SIMBA is an AI agent loop that mediates all interactions between the user and the analytical backend (**Figure 2a**). User queries, expressed in human language, are processed through a LLM Application Programming Interface (API), which interprets intent, selects appropriate tools, and generates executable configurations. These configurations are dynamically merged with system state and file context before triggering one or more analysis modules in the Python backend. Results are then collected, summarized, and returned to the user, forming a closed loop that supports iterative refinement and interactive analysis. This design enables users to specify high-level analytical goals while delegating procedural complexity to the system.

**Figure 2.**
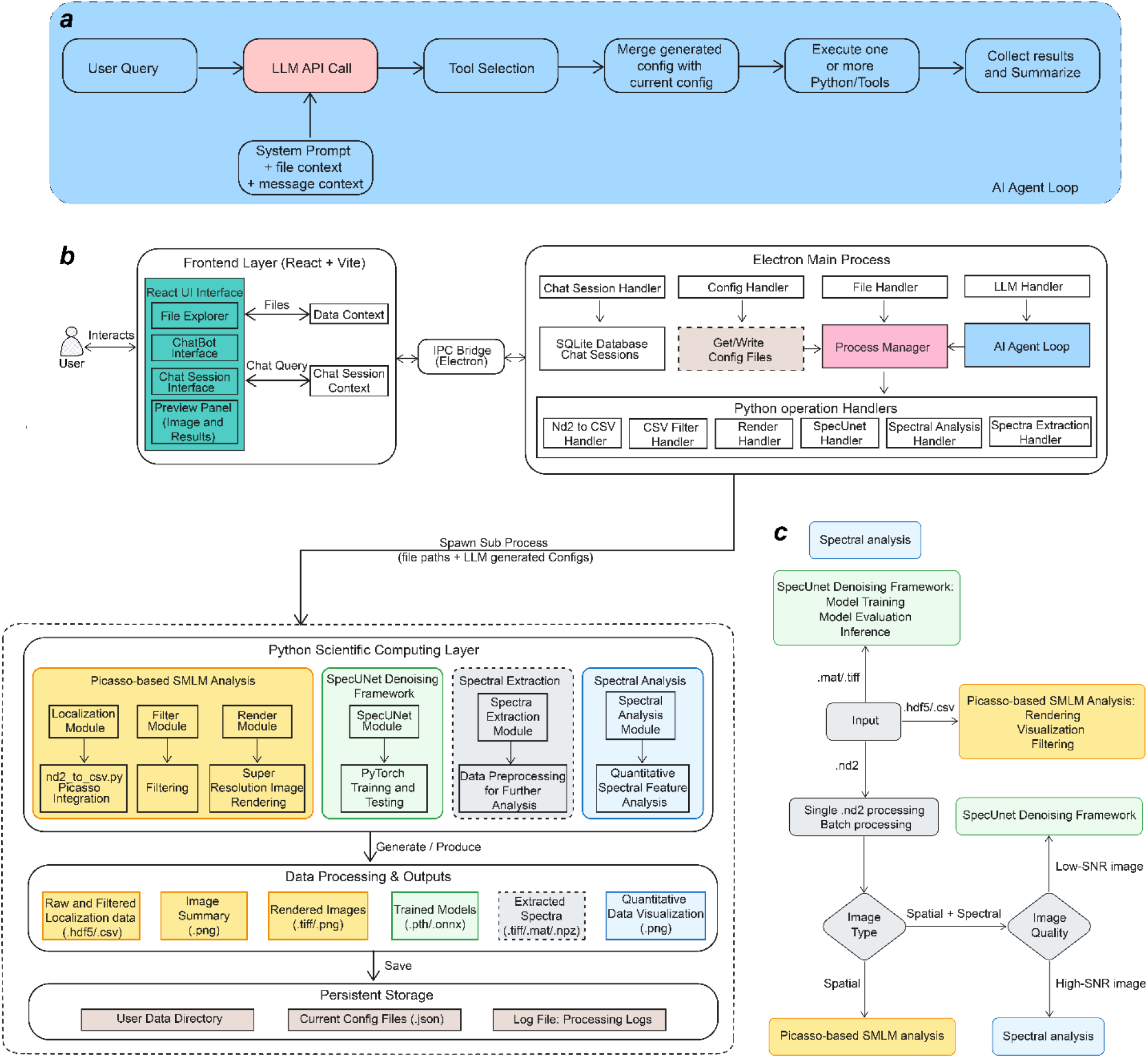
System architecture and domain specific workflow of the SIMBA desktop application. (***a***) The overall design of an agentic loop to develop AI agent to invoke, illustrating the extensible architecture designed to support scientific data analysis and visualization workflows; (***b***) Top left: frontend for file management, chat interaction, and visualization; top right: an Electron main process coordinating IPC communication, file operations, and frontend backend integration; bottom: a Python scientific computing layer for localization analysis spectral analysis, and filtering; and an AI agent loop that interprets user queries via LLM APIs, performs tool selection, integrates contextual configurations, executes analysis modules, and returns results; (***c***) Domain specific md-SMLM workflow illustrating a unified pipeline supporting multiple input formats, both single and batch processing modes, with adaptive routing based on modality and image quality.

The implementation of this agent loop is embedded within a layered software architecture that separates interface, orchestration, and computation (**Figure 2b**). The frontend layer (top left box in **Figure 2b**), built with React^50^and Vite^51^, provides an integrated user interface for file exploration, chat based interaction, and visualization of intermediate and final results. It includes dedicated components for dataset browsing, interactive plotting, and workflow feedback, allowing users to monitor execution and inspect outputs in real time. The Electron main process (top right box in **Figure 2b**) serves as the coordination layer for communication between the frontend, LLM services, local file systems, and backend computation modules. Inter-process communication (IPC) is used to transmit user queries, file metadata, and structured tool requests. Within this layer, specialized handlers manage chat sessions, configuration states, and LLM interactions, while also supporting prompt construction and response parsing. A process manager schedules and monitors computational tasks, enabling asynchronous execution and preventing interface blocking during long-running analyses. This layer also preserves persistent system state, such as user sessions, configuration histories, and execution logs. On the ohther hand, the backend Python scientific computing layer (bottom box in **Figure 2b**) contains modular analysis components, including (1) single-molecule localization directly process Nikon nd2 files or tiff files to localization table in a .csv format by reconfiguratng existing localization software such as Picasso^30^ to be excuative by agentic interaction and chat sessions, spectral extraction and analysis, filtering, rendering, and deep learning-based denoising using our recentl developed SpecUNet DL denoising framework.^27^ Each module is encapsulated with standardized input-output interfaces and exposed as a callable tool, allowing dynamic invocation by the orchestration agent. Execution is carried out through isolated subprocesses with structured parameter passing, which improves robustness and reproducibility while supporting task monitoring and cancellation. In short, these layers form a tightly integrated system in which user interaction, agent reasoning, and scientific computation are decoupled but seamlessly coordinated.

A key feature of SIMBA is the tight coupling between agent level reasoning and domain specific workflows (**Figure 2c**). The system supports multiple input formats, including .nd2, .hdf5, .csv, .mat, and .tiff, which are typically used in SMLM domain as well as single or batch processing of these filetypes. Upon ingestion, data are automatically routed based on modality and image quality. Spatial only datasets are processed through Python-based localization, rendering, and filtering pipelines, whereas datasets containing both spatial and spectral information are directed to spectral extraction and quantitative analysis modules. For low signal to noise conditions, an optional SpecUNet^27^ denoising step recently developed by us, can be invoked prior to spectral analysis, improving signal fidelity and downstream quantification. This adaptive routing mechanism allows SIMBA to generalize across diverse experimental conditions while maintaining analytical rigor.

The Python layer produces a range of outputs, including raw and filtered localization tables, reconstructed super resolution images, extracted spectra and statistical summaries, trained deep learning models, and quantitative visualizations. These outputs are persistently stored with user data, configuration states, and processing logs, ensuring reproducibility and traceability of all analytical steps. By integrating system level architecture with domain aware decision making, SIMBA transforms md-SMLM analysis from a fragmented sequence of manual operations into a cohesive, automated, and extensible computational framework.

### Multi-functional modules in SIMBA

To establish an end-to-end agent driven workflow for conventional SMLM analysis, we first implemented a fully integrated localization module within SIMBA that combines established algorithms with interactive system control (**Figure 3a**). This module builds upon Picasso^30^-based localization to process raw microscopy data, including molecule detection, filtering, and super resolution rendering, while embedding these steps within a unified and automated pipeline. A key feature of this implementation is the tight integration between parameter configuration and agent based execution. Users can directly specify localization parameters through a structured interface (**Figure 3b**), including fitting method, gradient threshold, sensitivity, and camera calibration settings. In parallel, the agent interface enables natural language driven control of the same workflow (**Figure 3c**), allowing users to initiate analysis, apply filters, and refine results without manual scripting. This dual mode interaction supports both expert level customization and accessible operation for non specialists.

**Figure 3.**
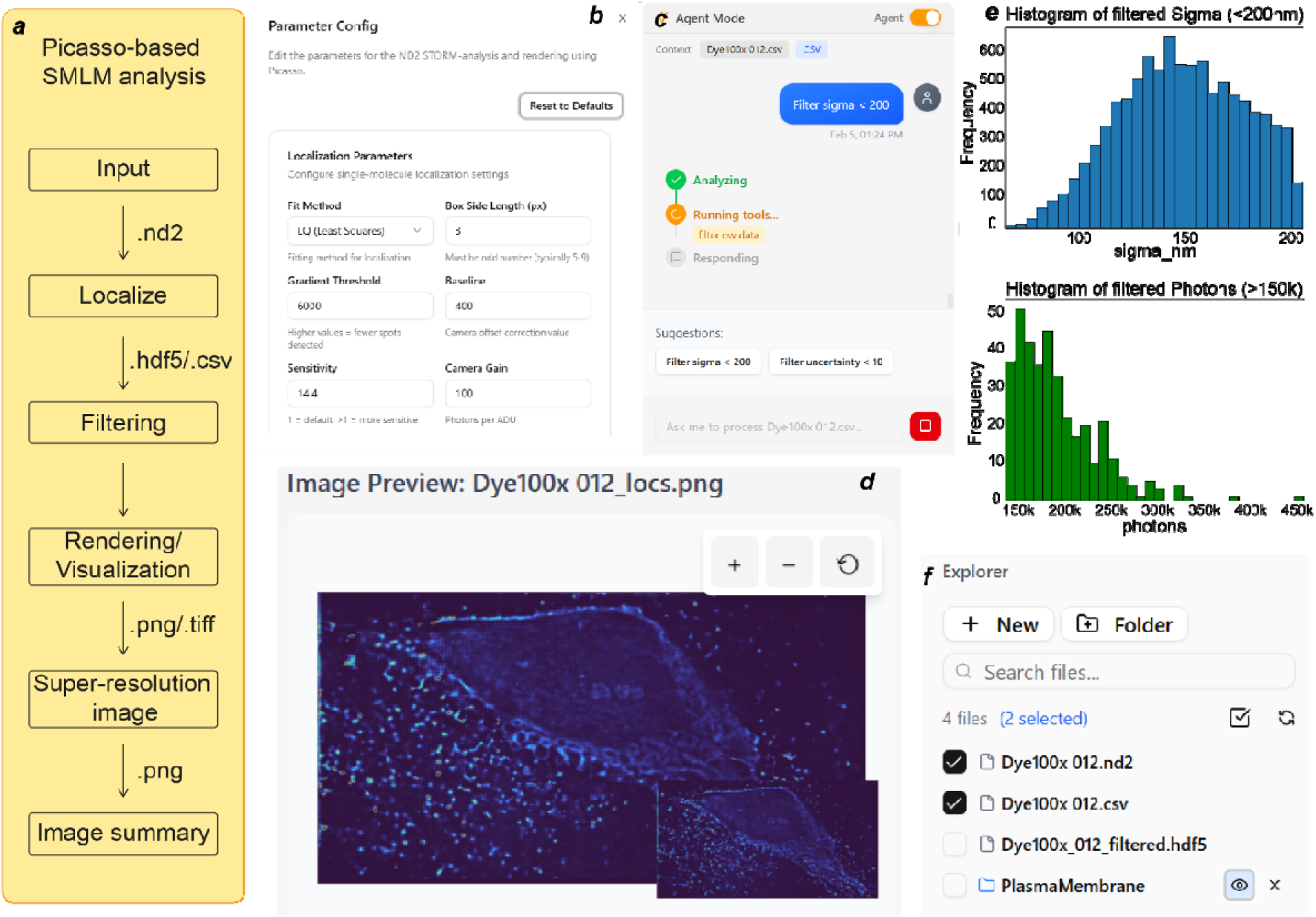
Agentic SMLM localization workflow and SIMBA interface. **(a**) Schematic of the Picasso based SMLM analysis pipeline implemented in SIMBA, including data input (.nd2), localization, filtering, rendering, and super resolution image generation. (**b**) Parameter configuration interface for localization analysis, allowing adjustment of fitting method, gradient threshold, sensitivity, and camera parameters. (c) Agent interaction interface showin natural language driven execution of analysis tasks and system status updates during processing. (d) Image preview panel displaying reconstructed super resolution images with interactive zoom and inspection. (e) Representativ statistical outputs of localization analysis, including distributions of localization precision (sigma) and photon counts after filtering. (f) File explorer for dataset management, enabling selection and organization of raw and processe files.

The system provides immediate visual feedback through an integrated preview panel (**Figure 3d**), where reconstructed SMLM images can be inspected in real time. This capability enables rapid validation of parameter choices and iterative refinement of analysis, addressing a common limitation in conventional workflows where feedback is delayed or fragmented across tools. Beyond visualization, SIMBA incorporates built in statistical analysis to support quantitative evaluation of localization results. As shown in **Figure 3e**, distributions of localization precision and photon counts are automatically generated following filtering steps, providing insight into data quality and enabling data driven parameter tuning. These outputs are directly linked to user actions, allowing the agent to recommend or apply additional filtering criteria based on observed distributions. The integration of file management within the same environment further streamlines the workflow (**Figure 3f**), enabling users to seamlessly navigate between raw datasets, intermediate outputs, and final results. This eliminates the need for external file handling and reduces the cognitive overhead associated with multi software pipelines. Automatic batch processing can also be achieved directly by agentic interactions and chat.

Building on the unified system architecture, SIMBA further implements domain specific analytical modules for spectral reconstruction and DL based denoising, both of which are seamlessly integrated within the agent driven workflow (**Figure 4**). These modules extend the capabilities of md-SMLM by enabling robust spectral quantification and improved signal recovery under challenging imaging conditions. The spectral analysis pipeline in SIMBA is designed as a structured yet flexible workflow that transforms raw imaging data into quantitative spectral features (**Figure 4a**). Input data in formats such as .nd2, .hdf5, or .csv are first subjected to background subtraction, followed by spectral extraction and calibration to generate single molecule spectra. These spectra are then analyzed to produce quantitative outputs, including spectral centroids, photon distributions, and wavelength dependent intensity profiles. Through the agent interface (**Figure 4b**), users can initiate and monitor spectral analysis using natural language commands, while underlying parameters are automatically configured or refined based on system context. Advanced users retain the ability to directly adjust parameters through dedicated configuration panels (**Figure 4c**), enabling precise control over spatial localization alignment and spectral cropping.

**Figure 4.**
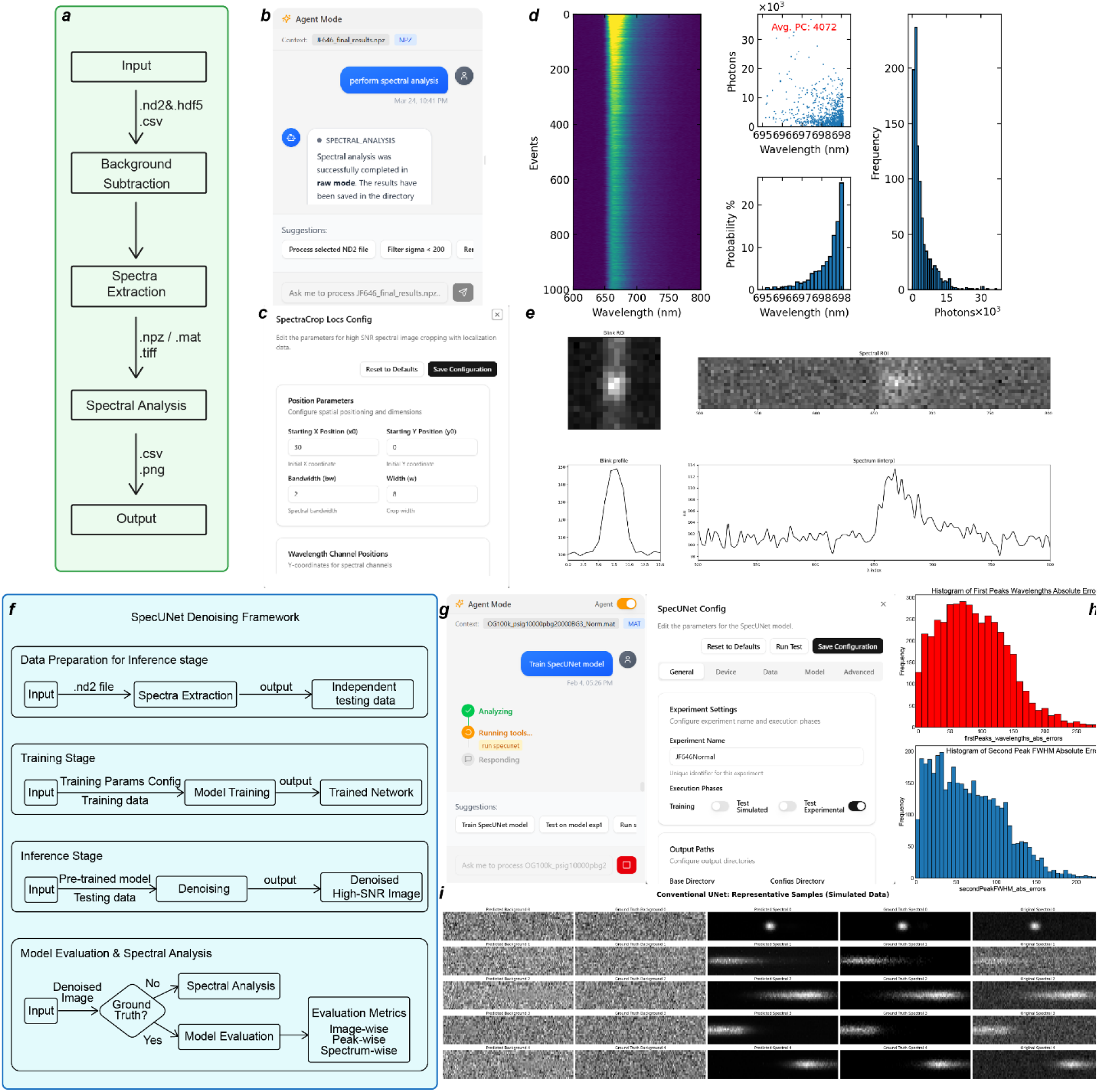
Agentic spectral analysis and AI denoising modules in SIMBA. (***a***) Schematic of the agentic spectral analysis workflow from input data to spectral reconstruction, including spectral extraction, calibration, and single molecule signal reconstruction; (***b***) SIMBA interface for spectral analysis, showing parameter configuration (***c***) and agent driven execution; (***d***) Representative outputs of spectral extraction with reconstructed single molecule spectra (left); and statistical analysis of spectral centroids and photon distributions across detected molecules (right); (***e***) raw single-molecule spatial and spectral images extracted on the live; (***f***) Schematic of the SpecUnet based denoising workflow, including data preparation, model training, inference, and evaluation; (***g***) SIMBA interface for DL model configuration and execution; (***h-i***) Representative comparison of raw and denoised spectral images (***i***), demonstrating signal recovery under low signal to noise conditions and (***h***) Quantitative evaluation of denoising performance, including error distributions of spectral peak position and full width at half maximum.

Representative outputs of the spectral module demonstrate both qualitative and quantitative capabilities (**Figures 4d-e**). Individual molecule spectra can be reconstructed from raw spectral images, while population level analyses reveal distributions of emission wavelength and photon counts across thousands of detected events. These outputs provide a direct link between imaging data and molecular level interpretation, supporting applications such as fluorophore identification and nanoenvironment sensing. To address the challenges of low signal-to-noise conditions inherent in md-SMLM, SIMBA incorporates a deep learning based denoising module based on the supervised learning SpecUNet framework (**Figure 4f**). This module follows a three stage pipeline consisting of data preparation, model training, and inference. During training, simulated or experimental datasets are used to optimize network parameters, producing a trained model that can subsequently be applied to unseen data. The agent interface enables users to configure and execute training or inference workflows with minimal manual intervention (**Figure 4g**), including selection of datasets, model parameters, and execution modes. The impact of denoising is illustrated through representative comparisons of raw and reconstructed spectral images (**Figure 4i**), where the SpecUnet model enhances signal clarity while preserving structural and spectral features. Quantitative evaluation based on experiment-informed monte-carlo simulations^27^ (**Figure 4h**) further demonstrates improved accuracy in key metrics such as spectral peak position and full width at half maximum, indicating enhanced precision in downstream spectral analysis. By integrating denoising directly within the analytical pipeline, SIMBA enables reliable extraction of spectral information even under low photon count conditions.

In summary, these modules highlight how SIMBA combines agent driven interaction with domain specific computation to enable end to end md-SMLM analysis. By unifying spectral reconstruction and deep learning based enhancement within a single framework, the system supports both high fidelity data interpretation and scalable analysis across diverse experimental scenarios.

## Discussions

SIMBA represents a shift from static and fragmented analysis pipelines toward dynamic, agent driven scientific workflows. By integrating localization, spectral analysis, and AI-based reasoning within a unified framework, SIMBA enables multidimensional single molecule imaging with reduced operational complexity and improved reproducibility. Importantly, this work reframes analysis not as a sequence of predefined steps, but as an adaptive process in which computational decisions evolve in response to data and user intent. A central advance is the introduction of agentic AI into super-resolution microscopy analysis. In contrast to conventional automation, which relies on fixed parameter settings and rigid pipelines, the SIMBA agent can interpret high-level user goals, select appropriate tools, and iteratively refine analysis strategies. This capability reduces dependence on expert knowledge while preserving analytical rigor, thereby narrowing the gap between accessibility and performance. Beyond usability, the agent framework also introduces a new layer of consistency by standardizing decision pathways that are otherwise highly user dependent.

Within the broader context of md-SMLM, SIMBA demonstrates how md-SMLM can be embedded into an extensible analytical paradigm. In the md-SMLM implementation, the platform enables simultaneous mapping of molecular identity and nanoenvironment, allowing spectrally overlapping fluorophores to be distinguished while quantifying local heterogeneity at the single molecule level. These capabilities support more reliable multiplexed imaging and provide access to information that is typically obscured in ensemble measurements, particularly in heterogeneous biological or material systems. Despite these advances, several challenges remain. The accuracy of spectral reconstruction is fundamentally constrained by photon budget and optical design, requiring careful balancing of spatial and spectral precision. In addition, while agent driven workflows improve adaptability, their reliability depends on well defined tool interfaces, robust error handling, and transparent decision making. Failure modes, particularly under low signal or ambiguous conditions, must be systematically characterized to ensure reproducibility across datasets and experimental settings. Future work will focus on strengthening both the analytical and methodological foundations of the platform. On the computational side, integrating advanced deep learning approaches for spectral denoising, feature extraction, and classification may further enhance performance under low signal conditions. On the systems level, expanding the tool ecosystem and incorporating additional dimensions such as fluorescence lifetime or polarization will extend the framework toward fully realized md-SMLM. Equally important is rigorous validation across diverse samples, imaging conditions, and instrument configurations to establish generalizability. Taken together, the integration of intelligent agents with multidimensional single molecule imaging introduces a new paradigm for microscopy analysis, in which flexibility, scalability, and interpretability are co designed. This approach has the potential to not only streamline existing workflows, but also enable new classes of measurements that were previously limited by analytical complexity.

## Methods

### md-SMLM optical setup

The single-molecule emission signals are collected by a fluorescence microscope equipped with a high-magnification objective (e.g., 100× Total Internal Reflection Fluorescence/TIRF) and passed through an imaging spectrometer to simultaneously capture the spatial and spectral images of individual molecules. A transmission grating is placed between a slit that is placed at an intermediate imaging plane and a pair of relay lenses (f = 100 mm) to form an imaging spectrometer. The imaging spectrometer predominantly splits the photons between the 0th- and 1st-order diffraction in a ratio of about 20:80 in the spectral range of 500-800 nm. The 1st order single-molecule emission is chromatically dispersed to encode wavelength-dependent intensities in the horizontal axis of a scientific Complementary Metal–Oxide–Semiconductor (sCMOS) camera. The position of the grating along the optical axis can be adjusted to tune the spectral dispersion from 1.5 to 5 nm/pixel. The detailed view of a representative single-molecule spectral image depicted poor SNR.

### PAINT imaging acqusition

Typical PAINT imaging was performed based on the aforementioned md-SMLM setup. The entire imaging spectrometer was replaced by an Electron Multiplying Charge-Coupled Device (EMCCD) camera (iXon Ultra 897) which was placed at the intermediate imaging plane where the slit was originally located to achieve conventional SMLM imaging capabilities. After the cells were incubated with 2 (1 μM) in a mixture of phosphate-buffered saline (PBS) and dimethyl Sulfoxide (DMSO) (100:1, v/v) for ∼30 min, they were immediately used for super-resolution imaging.

### Sample preparation

Methanol solutions of JF646 (1 nM, 200 μL, a gift from Dr. Luke Lavis’s lab, Janelia Research Campus) were spin-coated onto bare glass substrates (#1 premium cover glass, 22 × 22 mm^2^, Fisher) using an EZ4 programmable Spin Coater for 1 min at 2000 rpm, dried under vacuum, and mounted onto the microscope stage to be imaged on the same day. A drop of antifade mounting medium was added to the fluorescence nanosphere sample and sealed onto a cover slide. For cell imaging, the cells prepared after cell culture were washed with PBS, transferred to a stage-top live-cell incubator (Okolab), and mounted on the microscope for imaging directly at 37 °C with CO2 (5%). For cholesterol addition, the cells were incubated with 10-μM cholesterol (Sigma C4591) before staining. For the cholesterol-treated image, the cells were incubated with (compound?)2 (1 μM) in a mixture of phosphate-buffered saline (PBS) and dimethyl sulfoxide (DMSO) (100:1, v/v), and they were immediately used for super-resolution imaging.

### Imaging acquisition and processing

For HT-SMS experiments of JF646, single-molecule images were acquired with a camera exposure time of 20 ms. The spectroscopic response of the imaging spectrometer was calibrated with a fluorescence lamp before each imaging session, and the spectral dispersion was 2.355 nm/pixel. The entrance slit width was adjusted to 50 μm, and the microscope stage was automatically scanned at a speed of ∼5 μm s□^1^. JF646 samples were excited at 640 nm with a power density of 14 W cm□^2^. For PAINT imaging, images were acquired using a Changchun New Industries Optoelectronics MGL-FN-589 continuous-wave laser operating at 589 nm with a power density of 330 W cm□^2^, with an exposure time of 10 ms and 50,000 frames collected per acquisition.

### Cell Culture

Human bone osteosarcoma epithelial cells (U2OS line, ATCC HTB-96) were grown in McCoy’s 5A medium supplemented with fetal bovine serum (10%, FBS, Fisherbrand™), penicillin/streptomycin (1%,100 U mL–1, GibcoTM) at 37 °C with CO2 (5%). The cells were seeded in an 8-well chambered cover glass (Cellvis, C8SB-1.5H) with 50–70% confluency. After 24 hours of seeding, the cells were proceeded with dye staining.

### Agentic AI framework design and tool orchestration

SIMBA uses an agentic framework in which an LLM functions as an orchestration layer rather than a direct generator of scientific results. Upon receiving a natural language-based query, the LLM interprets the user’s request, selects the correct tool for analysis from a predefined tool set, and maps the user’s request to structured parameters, finally executing it through the backend. The tools are predefined based on structured JSON schemas, which are compatible with function calling protocols, including microscopy image preview, localization conversion, batch processing, localization rendering, CSV filtering, plotting, spectra extraction, spectral analysis, configuration retrieval, file-context inspection, and SpecUNet-based denoising. Metadata from user-selected files is provided to the LLM together with the query to determine input compatibility, identify the appropriate workflow, and generate follow-up actions. Once the tool is identified, the backend will validate the input and merge the user-defined configuration before executing the task through the corresponding backend handler. The results are summarized in natural language and then sent back to the interface, completing a loop of interpretation, execution, and response that enables multi-step microscopy workflows through a single conversational interface while ensuring interpretability and reproducibility.

### SIMBA system architecture

SIMBA was implemented as a hybrid desktop software system that integrates LLM-based natural-language interaction with validated scientific image analysis modules. The system adopts a function-calling architecture in which the LLM serves as an orchestration layer that interprets user requests, selects appropriate analysis tools, extracts relevant parameters, and coordinates execution, At the same time, all scientific computation is performed using predefined domain-specific algorithms. The overall software stack comprised a React- and TypeScript-based frontend, an Electron layer served as an intermediate coordination layer, and Python-based scientific computing backend modules for localization, denoising, and spectral analysis

### Frontend

The frontend layer was developed using React^50^ and Vite^51^, with Tailwind CSS^52^ and Shadcn/Radix^53^-based components for the desktop user interface and React-Plotly.js for interactive scientific visualization. The frontend provides integrated interfaces for file exploration, chat-based interaction, configuration editing, and visualization of intermediate and final outputs. A hooks-based architecture was used to organize agent interaction and execution state. In particular, the useAgentChat workflow manages conversation state and tracks execution phases such as thinking, tool execution, and response generation, whereas the useAgentExecution. The workflow maintains a registry of callable functions corresponding to the available tools. Runtime parameters are resolved by merging explicit user-provided inputs with saved configuration defaults, allowing both flexible user control and reproducible execution settings.

### Electron layer

The desktop application layer was implemented with Electron and Electron IPC to mediate communication between the user interface, the LLM service, the file system, and the local Python environment. Within this layer, dedicated services handle chat session persistence, configuration management, file operations, and LLM requests, while a process manager oversees execution of computational tasks. Local persistence of chat sessions and application state was supported by SQLite.

### Python backend

The scientific computing backend consisted of modular Python components for microscopy data analysis. These modules included Picasso-based localization for converting raw ND2 microscopy data into localization tables, rendering and filtering utilities for downstream super-resolution analysis, spectral extraction workflows serve as a pre-processing function to extract single molecule spectra from md-SMLM images for downstream analysis, and spectral analysis for downstream wavelength-based quantification. SIMBA also integrated a SpecUNet-based deep learning module for spectral image denoising, with support for training on simulated data and testing on both simulated and experimental datasets. Scientific analysis modules were implemented in Python using packages including Picasso, PyTorch, NumPy, Pandas, SciPy, h5py, hdf5storage, tifffile, nd2, Pillow, Matplotlib, and imageio. Each module was exposed in a standardized parameter interface so that it could be invoked programmatically from the Electron layer.

### Data flow

During execution, user queries entered through the frontend were combined with the currently selected file context and forwarded through Electron IPC to the backend. The backend then assembled the corresponding LLM request and tool definitions, received structured tool calls, validated inputs, and launched the appropriate backend handler or Python subprocess. Python modules were invoked as subprocesses with JSON-based parameter passing, providing process isolation, explicit control of inputs and outputs, and compatibility with cancellation and logging. Resulting outputs, including processed data files, images, and analysis summaries, were returned through the backend to the frontend for visualization and user feedback. This implementation enabled SIMBA to support iterative and conversational microscopy workflows while maintaining consistent configuration-aware execution.

### Implementation of SpecUNet

SpecUNet was implemented in MATLAB and Python and trained on a personal computer equipped with a single NVIDIA GeForce RTX 4070 or 3080 Ti GPU. The effects of sample size and key hyperparameters, including learning rate and number of training epochs, were evaluated during model optimization. Training and validation datasets were divided at a ratio of 80:20, and an independent test dataset was generated separately under the same simulation conditions. Detailed descriptions of the network architecture, convolutional filter sizes and additional training settings are provided in Note S2 of the previous study.^Ref^ For integration into SIMBA, the original SpecUNet codebase was translated into Python and reorganized as a standard installable package 2. The package was built as a distributable wheel using `python -m build`, allowing installation either from a package repository or from a locally provided wheel file. After installation through `pip`, SpecUNet became available within the SIMBA analysis environment [Future Python codebase citation]. Upon user request, SIMBA invokes the packaged SpecUNet module and executes the corresponding training or inference functions, enabling both training and testing on newly simulated md-SMLM datasets and denoising of preprocessed experimental datasets during inference.

## Data Availability

SIMBA desktop application, tutorials, and example data are available at https://simba-website-indol.vercel.app for testing; raw imaging data are available upon reasonal request to the corresponding author.

## Acknowledgements

We acknowledge the funding support from the National Science Foundation (CHE-2246548 and CHE-2441081), the National Institutes of Health (R35GM155241 and R35GM156520), the North Carolina State University Comparatie Medicine Institute Award and the College of Engineering Applied AI Research Accelerator Award.

## Competing interests

The authors declare no competing interests

## Author contributions

Y.Z., D.X., C.L. conceptualized and supervised the projects; H.Mao. led the overall SIMBA development and experimental studies, H.Mauny contributed to the front-end development, O.K.V. contributed to the back-end development; H.Mao and Y.Z. drafted the manuscript and all the remaining authors contributed to manuscript editing.

## Notes

https://simba-website-indol.vercel.app/

### Competing Interest Statement

The authors have declared no competing interest.

